# Self-organized stem cell-derived human lung buds with proximo-distal patterning and novel targets of SARS-CoV-2

**DOI:** 10.1101/2021.01.06.425622

**Authors:** E.A. Rosado-Olivieri, B. Razooky, H.-H. Hoffmann, R. De Santis, C.M. Rice, A.H Brivanlou

**Author notes:** These authors contributed equally to this work (E.A. Rosado-Olivieri, B. Razooky).

## Abstract

Severe acute respiratory syndrome coronavirus 2 (SARS-CoV-2) is the causative agent of the global COVID-19 pandemic and the lack of therapeutics hinders pandemic control^1–2^. Although lung disease is the primary clinical outcome in COVID-19 patients^1–3^, how SARS-CoV-2 induces tissue pathology in the lung remains elusive. Here we describe a high-throughput platform to generate tens of thousands of self-organizing, nearly identical, and genetically matched human lung buds derived from human pluripotent stem cells (hPSCs) cultured on micropatterned substrates. Strikingly, *in vitro*-derived human lung buds resemble fetal human lung tissue and display *in vivo*-like proximo-distal coordination of alveolar and airway tissue differentiation whose 3D epithelial self-organization is directed by the levels of KGF. Single-cell transcriptomics unveiled the cellular identities of airway and alveolar tissue and the differentiation of WNT^hi^ cycling alveolar stem cells, a human-specific lung cell type^4^. These synthetic human lung buds are susceptible to infection by SARS-CoV-2 and endemic coronaviruses and can be used to track cell type-dependent susceptibilities to infection, intercellular transmission and cytopathology in airway and alveolar tissue in individual lung buds. Interestingly, we detected an increased susceptibility to infection in alveolar cells and identified cycling alveolar stem cells as targets of SARS-CoV-2. We used this platform to test neutralizing antibodies isolated from convalescent plasma that efficiently blocked SARS-CoV-2 infection and intercellular transmission. Our platform offers unlimited, rapid and scalable access to disease-relevant lung tissue that recapitulate key hallmarks of human lung development and can be used to track SARS-CoV-2 infection and identify candidate therapeutics for COVID-19.

---

The emergence of SARS-CoV-2 in late 2019 sparked an explosive global pandemic of COVID-19 disease, with > 75 million confirmed cases and > 1.6 million deaths to date^1–2^. The rational design of COVID-19 therapies will require an understanding of the lifecycle of the virus during infection in human cells. Key infection routes of SARS-CoV-2 involve the nasal passages, lung airways and alveoli^1–3^. In particular, the lung is the one of the most vulnerable target organs for SARS-CoV-2 as acute lung injury and pneumonia-associated complications are primary clinical outcomes in severe cases of COVID-19^1–3^. In this organ, airway multi-ciliated cells and alveolar type 2 (AT2) pneumocytes are the primary targets of this virus as they express the receptor ACE2 as well as a myriad of factors that regulate viral entry into host cells^3,5–9^. How SARS-CoV-2 induces local tissue damage and pathology in the lungs remains elusive.

Current models of SARS-CoV-2 lung infection rely on the *in vitro* culture of human primary lung tissue^3,10–12^, which remains challenging due to its unpredictable quality as well as its high inter-donor phenotypic and genetic variability. These features are of particular importance as genetic heterogeneity plays a large role in SARS-CoV-2 replication and outcome^13^. To circumvent these challenges, directed differentiation protocols have been developed to differentiate human pluripotent stem cells (hPSCs) into lung airway and alveolar cells as an alternate source of tissue to study lung biology and disease (Jacob et al. 2017; McCauley et al. 2017; Dye et al. 2015; Miller et al. 2019). These models recapitulate human development by deploying a set of signals known to coordinate lung development *in vivo*^14–17^. More recently, hPSCs-derived lung cells were used to study cellular responses upon SARS-CoV-2 infection^18–19^. Similarly, an organoid-based platform for generating lung cells from hPSCs was recently used to identify small molecules that halt SARS-CoV-2 infection^18^. This highlights the potential of using hPSC-based platforms for the study of SARS-CoV-2-associated lung pathology and for high-throughput identification of therapeutics for COVID-19. Although these stem cell-based models offer an inexhaustible supply of human lung cells, they lack the controlled tissue organization observed in developing and adult lung tissue such as the coordinated segregation of alveolar and airway tissues^20–21^. In addition, current protocols take 1-3 months to differentiate lung cells from hPSCs underscoring a need to develop fast and scalable platforms to generate these cells *in vitro*.

## Reconstituting human lung development on confined geometries

To overcome these limitations, we sought to develop a micropattern-based platform to generate synthetic embryonic tissues that model fetal human lungs. This technology allows us to generate synthetic tissues that model the *in vivo* embryonic counterparts by exploiting the self-organizing capabilities of hPSCs when they are cultured on confined geometries in micropattern chips^22–23^. The generation of self-organized lung tissues relies on the stepwise modulation of signaling pathways that direct lung development *in vivo* (Fig. 1A)^14–17^. As lung progenitor cells are derived from anterior endodermal progenitors in the embryo, we first induce hPSCs to differentiate into SOX17+ definitive endoderm in standard monolayer cultures, by applying WNT and ACTIVIN stimulation for three days (Fig. 1B, Extended Data Fig. 1A-B). Endodermal cells were then seeded on micropattern substrates and exposed to the TGFβ inhibitor SB431542 (SB) and BMP inhibitor Dorsomorphin (DM) for another 3 days to induce FOXA2+ anterior endoderm (foregut; Fig. 1C, Extended Data Fig. 1A-B)^15–16^. Finally, cells were exposed to WNT-activation, KGF, BMP4, and retinoic acid (RA) stimulation for 7 days, to promote the differentiation of NKX2.1+ multipotent lung progenitors (Fig. 1D). These progenitors give rise to both airway and alveolar cell types^20–21^. We find that compared to standard monolayer differentiation protocols where KGF is dispensable^15–16^, a combined induction by KGF and BMP4 together, rather than each alone led to a significant increase in the proportion of NKX2.1+ cells on lung progenitor micropatterned colonies (Fig. 1E, Extended Data Fig. 1C-E). Lung progenitor fate depended on the levels of KGF and BMP4, as we detected a dose-dependent increase in the proportion of NKX2.1+ progenitors in micropatterned cultures (Fig. 1I-J, Extended Data Fig. 2A-C). This platform allows us to generate thousands of lung progenitor colonies of defined sizes in a single micropattern chip (Extended Data Fig. 1F).

**Figure 1:**
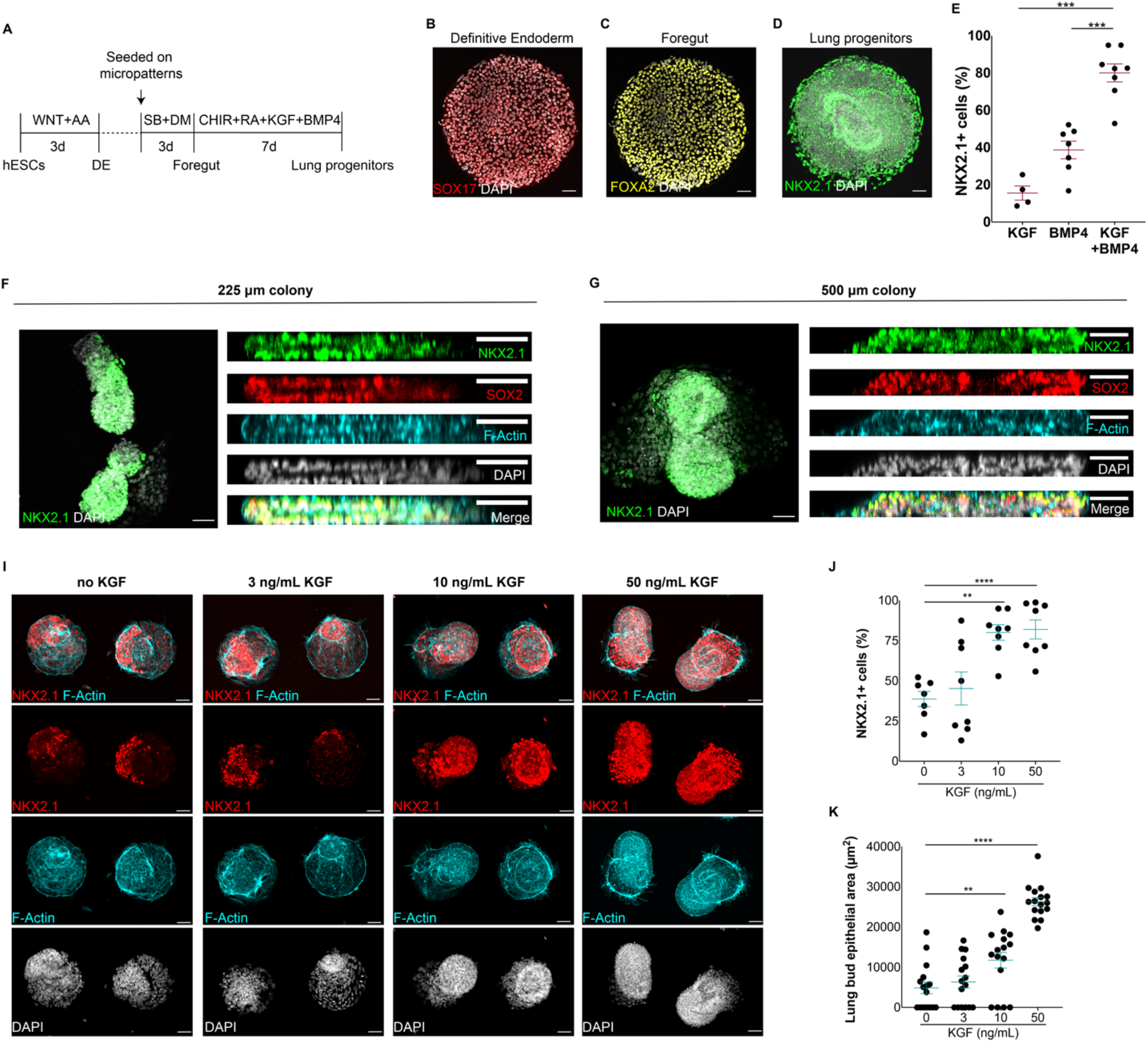
Generation of self-organized epithelial lung buds on confined geometries. A) Protocol for the generation of synthetic lung buds on confined geometries in micropatterns. B-D) Generation of SOX17+ endoderm cells (B), FOXA2+ anterior endoderm cells (C) and NKX2.1+ multipotent lung progenitors (D) at the end of definitive endoderm (DE), foregut and lung progenitor induction stages, respectively. E) Proportion of NKX2.1+ lung progenitors upon modulation with BMP4, KGF or BMP4/KGF for 7 days. F-G) Top and side views of 3D epithelial buds containing NKX2.1+ multipotent lung progenitors in 225 and 500 μm colonies. I) Micropattern colonies containing NKX2.1+ epithelial buds at increasing doses of KGF. J) Proportion of NKX2.1+ progenitor cells at increasing doses of KGF. K) Quantification of lung bud epithelial area in micropattern colonies at varying doses of KGF. (**p<0.01,***p<.001, ****p<0.0001, Dunnett’s multiple comparison test; scale bar, 50 μm)

During early human development *in vivo*, fetal lung progenitors arise in epithelial buds that form from an outpouching of the anterior endoderm, co-expressing NKX2.1 and SOX2, around week 4 of gestation^20–21^. Later, NKX2.1 expression becomes confined to the alveoli, while SOX2 expression specifically demarcates the airway^20–21^. Interestingly, we find that upon induction, multipotent lung progenitors self-organize into three-dimensional (3D) epithelial chords containing NKX2.1+/SOX2+ progenitors (Fig. 1F-G), reminiscent of fetal lung buds *in vivo*^20–21^. We observed lung progenitor epithelial buds on circular colonies of 225 to 800 μm of diameter with the number of individual buds forming restricted to only one in 225 μm colonies (Fig. 1F-G, Extended Data Fig. 1F). We find that the levels of KGF direct the epithelialization and self-organization of lung progenitors as there is a dose-dependent increase in the epithelial area of lung buds upon KGF modulation (Fig. 1I-K, Extended Data Fig. 2A). High doses of KGF (50 ng/mL), robustly induce epithelial lung buds that are nearly identical in size and morphology (Fig 1I-K, Extended Data Fig. 2A). Our model highlights the self-organizing capabilities of stem cell-derived lung progenitors on confined circular geometries which can be used to quantitatively dissect morphogenetic events in the embryonic human lungs.

## Proximo-distal patterning of fetal-like human lung buds

Upon induction of the lung primordium, epithelial buds are further patterned along their proximo-distal axis which leads to a coordinated segregation of proximal SOX2+ airway and distal NKX2.1+/SOX9+ alveolar progenitors^20–21^. This initial patterning event is critical for the coordination of region-specific tissue morphogenesis and the differentiation of airway and alveolar cells that will ensue^20–21^. These features that are not faithfully recapitulated in current lung organoid protocols^14–18^. Strikingly, lung progenitors grown on small confined geometries (225 μm diameter) display proximo-distal coordination of progenitor differentiation with a SOX9+ alveolar-like and a SOX2+ airway-like located in non-overlapping tissue domains (Fig. 2A-B), as in embryonic fetal lung buds^20–21^. SOX9+ alveolar-like progenitor cells are positioned proximal to the micropattern surface whereas SOX2+ airway-like progenitor cells are located more distally forming a continuous 3D epithelial structure (Fig. 2A-B). These synthetic human lung buds also display early hallmarks of cellular differentiation and contain airway multi-ciliated cells (AcTub+; Fig. 2C) as well as alveolar type 1 (HOPX+, Fig. 2D) and type 2 pneumocytes (proSFTPC+, Fig. 2C). In 500 μm lung progenitor colonies, we also detected airway basal stem cells (P63+) and mucus-producing goblet cells (MUC5AC+) in epithelial cords that were less organized compared to 225 μm micropatterns (Extended Data Fig. 3A-E). Thus, lung progenitor cells cultured on confined geometries self-organize into fetal-like human lung buds with proximal-distal coordination of tissue differentiation.

**Figure 2:**
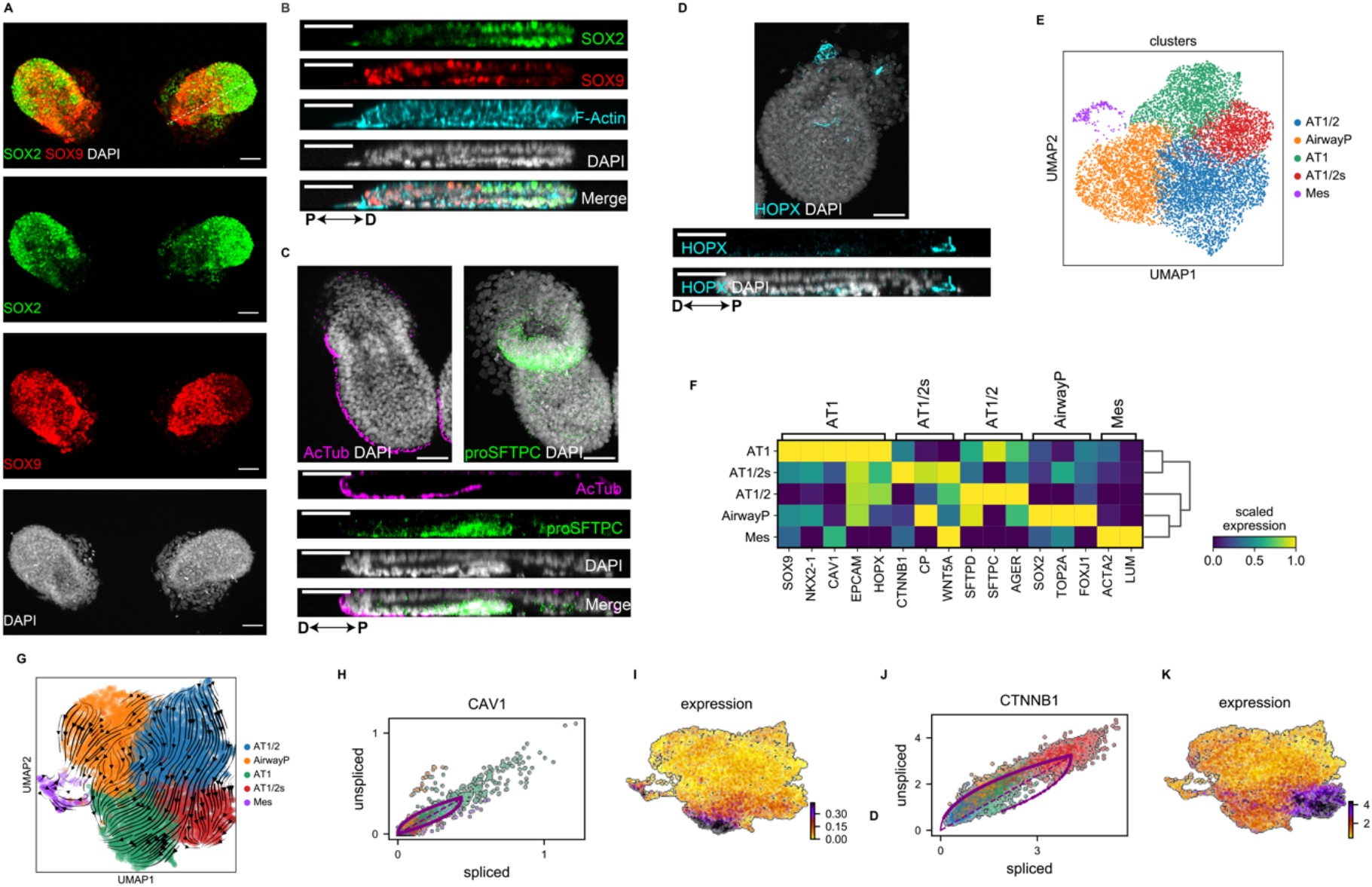
Proximo-distal coordination of airway and alveolar tissue differentiation in synthetic human lung buds. A-B) Top (A) and side (B) views of synthetic lung buds with proximo-distal segregation of SOX2+ airway and SOX9+ alveolar progenitors. C-D) Top view and side view of synthetic lung buds and identification of multi-ciliated (AcTub+, C), type 2 pneumocytes (proSFTPC+, C) and type 1 pneumocytes (HOPX+, D). E) UMAP plot and identification of 5 major cell clusters in synthetic lung buds by single cell RNA-sequencing. F) Heatmap of scaled gene expression levels in each of the identified cell types of cell type-specific lung markers of alveolar AT1 (CAV1+), AT1/2 (AGER+,SFTPD+), and AT1/2s (CTNNB1+), airway progenitors (SOX2+, FOXJ1+, TOP2A+) and mesenchymal cells (ACTA2+, LUM+). G) RNA velocity plots of proposed lineage trajectories (denoted by predicted arrows) among cell types in synthetic lung buds. H-K) Levels of unspliced and splice RNA transcripts (H, J) and gene expression (I, K) of the AT1 marker CAV1 (H-I) and AT1/2s marker CTNNB1 (J-K). Cells in H and J are colored according to the cluster classification in G. Scale bars: 50 μm. (proSFTPC: pro-surfactant protein C; SOX9+ Prog: SOX9+ progenitors; SOX2+ Prog: SOX2+ progenitors; Mes: mesenchyme; D: distal; P: proximal).

## Molecular signature of standardized human synthetic lungs

Single cell RNA-sequencing analysis of self-organizing human lung buds grown on circular micropatterns (225 μm diameter), identified 5 major cell subpopulations including SOX2+ airway progenitors (AirwayP), early differentiating alveolar cells (AT1/2), stem cell-like alveolar cells (AT1/2s), type 1 pneumocytes (AT1) as well as mesenchymal (Mes) cells (Fig. 2E-F, Extended Data Fig. 4A-E). SOX2+ airway cycling progenitors can be identified based on the high expression of SOX2 and FOXJ1 as well as TOP2A, a cell cycle marker (Fig. 2F, Extended Data Fig. 4D-E). Early AT1/2 cells can be identified based on the co-expression of the AT1 marker AGER as well as surfactant proteins SFTPD, SFTPC and SFTPB (Fig. 2F, Extended Data Fig. 4E). AT1 cells display high levels of SOX9 and NKX2-1 as well as the canonical AT1 markers CAV1 and HOPX (Fig. 2F, Extended Data Fig. 4D-E). Stem cell-like AT1/2s display high levels of the canonical AT2 marker ETV5 as well as CTNNB1, CP, WNT5A, TCF7L2 (Fig. 2F, Extended Data Fig. 5A), which are marker genes of a human-specific stem cell-like alveolar cell type recently identified in adult lung tissue^4^. Compared to other alveolar cell subpopulations, AT1/2s display high expression of the cell cycle marker TOP2A, suggesting that these cells correspond to cycling alveolar stem cells (Extended Data Fig. 5A). Interestingly, we also identified a population of stromal-like cells based on the expression of lumican (LUM), collagen (COL3A1, COL1A2), and ACTA2, a marker of lung mesenchyme (Fig. 2F, Extended Data Fig. 4D-E)^24–25^.

To ascertain lineage dynamics in synthetic lung buds, we performed RNA velocity analysis that predict lineage trajectories based on transcriptional dynamics and splicing kinetics among cells^26–27^. We identified three independent lineages including mesenchymal, airway, and alveolar differentiation trajectories (Fig. 2G). Among alveolar cells, our analysis suggests that early AT1/2 give rise to both AT1 and stem cell-like AT1/2s (Fig. 2G). We further resolved gene expression correlates of alveolar lineage bifurcation (Extended Data Fig. 5B-D) and detected hallmarks of alveolar cellular differentiation including an increase in CAV1 expression, a marker of AT1, along the AT1 differentiation trajectory, and CTNNB1 (β-catenin) along the AT1/2s differentiation trajectory (Fig. 2H-K). These results are consistent with recent findings that suggest that high WNT signaling is a key hallmark of stem cell-like alveolar cells^4^. Our analysis provides new insights on early differentiation events in fetal lung buds and resolves unprecedented developmental relationships among alveolar cells.

## Tracking SARS-CoV-2 infection and transmission in synthetic human lung buds

As our synthetic human lung buds contain the proposed cellular targets of SARS-CoV-2^3^ and endemic coronaviruses, we sought to determine whether we can track SARS-CoV-2 infection and pathology in these tissues. We observed viral tropism for endemic coronaviruses HCoV-229E, HCoV-OC43 and, to a lower extent, HCoV-NL63 (Extended Data Fig. 6), suggesting that human lung buds are a tractable model to study respiratory infection by human coronaviruses. To determine if our platform recapitulates COVID-19 pathology, we infected synthetic lung buds at different stages of lung bud formation with a patient-derived isolate of SARS-CoV-2 (USA-WA1/2020; Fig. 3A-C, Extended Data Fig. 7A-C). We detected viral tropism in both SOX2+ airway and SOX9+ alveolar cells (Fig. 3A-C, Extended Data Fig. 7A-C), including in multi-ciliated cells and type 2 pneumocytes (Figure 3G-H), primary targets of SARS-CoV-2^3^. In contrast to experiments with primary lung tissue cultured *in vitro*^3^, we identified an increased susceptibility to infection in alveolar cells compared to airway cells (Fig. 3C) and identified type 1 pneumocytes (HOPX+) as additional targets of SARS-CoV-2 (Fig. 3I). Consistent with these results, we detected higher expression of the receptor ACE2 as well as proteases TMPRSS2 and FURIN, which regulate SARS-CoV-2 entry into target cell types^1,28–29^, in alveolar cell types compared to airway cells (Extended Data Fig. 5E-F).

**Figure 3:**
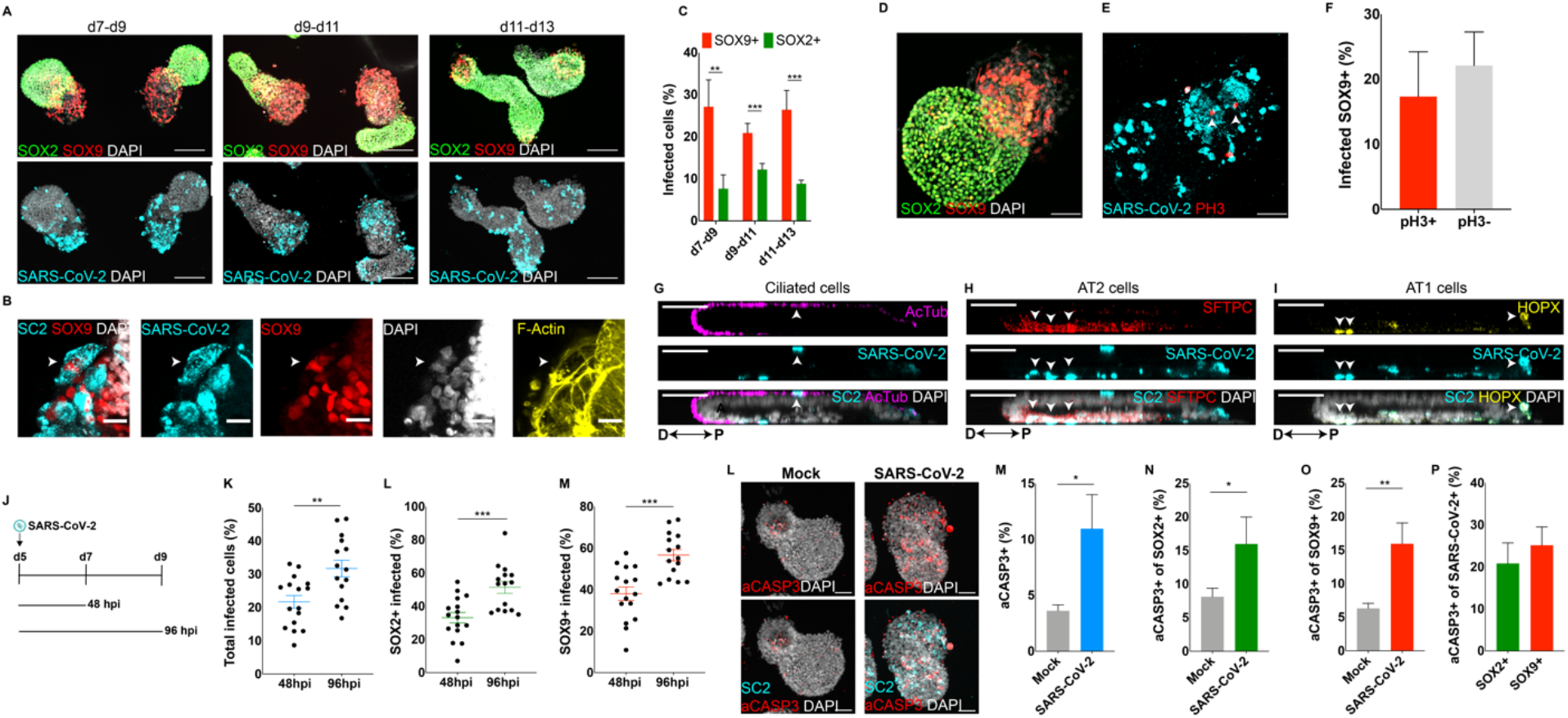
SARS-CoV-2 infects alveolar and airway epithelial cells in synthetic human lung buds. A) Synthetic lung buds infected with SARS-CoV-2 at day 7, 9 or 11 of lung bud formation and collected 48 hours post-infection (hpi) (scale bar: 100 μm). B) SARS-CoV-2 infection in SOX9+ alveolar cells (scale bar: 50 μm). C) Percentage of infected cells in SOX9+ alveolar and SOX2+ airway cells. D-F) Detection and quantification of SARS-CoV-2-infected pH3+ cycling SOX9+ alveolar cells. Arrows denote infected pH3+ cycling cells. (scale bar: 50 μm) G-I) Detection of SARS-CoV-2 infection in multi-ciliated (AcTub+; G), type 2 pneumocytes (proSFTPC+; H) and type 1 pneumocytes (HOPX+; I). Arrows denote infected cells expressing cell type-specific markers (scale bar: 50 μm). J) Experimental scheme to track SARS-CoV-2 transmission in synthetic lung buds. K-M) Percentage of infected cells of total, SOX2+ and SOX9+ cells 48hpi and 96hpi. N-L) Detection and quantification of active CASP3+ apoptotic cells in mock and SARS-CoV-2 infected synthetic lung buds (scale bar: 50 μm). (*p<0.05, **p<0.01,***p<.001, ****p<0.0001, Dunnett’s multiple comparison test; SC2: SARS-CoV-2; aCASP3: active Caspase-3; PH3: phospho-Histone H3; D: distal; P: proximal)

Interestingly, we identified tropism in phospho-Histone3 (pH3)-positive cycling SOX9+ alveolar cells, which correspond to AT1/2s, consistent with the high expression of ACE2 and related proteases observed in this cell type (Fig. 3D-F, Extended Data Fig. 5E-F). We did not observe significant differences in the levels of infection between dividing and non-dividing SOX9+ alveolar cells (Fig. 3F). Our analysis suggests that cycling alveolar stem cells may also represent targets of SARS-CoV-2 in alveoli, with important clinical implications for COVID19.

Following a viral transmission paradigm in which we tracked infection 48- and 96-hours post-infection (Fig. 3J), we observed an increase in the number of infected cells over time in a cell type-independent manner (Fig. 3K-M). This suggests that intercellular transmission (viral spread) was occurring in synthetic human lung buds. We also observed high levels of apoptosis upon infection in both alveolar and airway tissues compared to mock-infected synthetic lung buds mimicking virus-induced lung pathogenesis (Fig. 3L-P). Using this model, we can recapitulate the entire viral lifecycle and are now able to track cell type-dependent susceptibilities to infection, intercellular transmission, and cytopathic effects in hundreds of individual synthetic lung buds grown in a single micropattern chip.

## Synthetic human lungs for comparative therapeutics screens

As our platform mimics *in vivo* features of lung tissue and is scalable, we next screened therapeutics that were shown to limit SARS-CoV-2 infection through binding assays and in cell lines^30^. We tested human antibodies isolated from convalescent plasma of patients^30^ and evaluated the efficiency of the antibodies in neutralizing SARS-CoV-2 and preventing infection (Fig. 4A). Each neutralizing antibody, nAb, compared to a control (Fig. 4B), was tested at a range of dilutions (Fig. 4C-E). We found that each potently inhibited SARS-CoV-2 infection (Fig. 4C-E) at IC50s similar to those found for cell lines^30^. The nAbs also seemed to broadly inhibit SARS-CoV-2 infection, rather than in a cell-type specific manner (Fig. 4F-G). As the initial assay shows that these nAbs can inhibit infection when incubated with the virus prior to infection, we next wanted to test how pragmatic therapeutic approaches where the nAb is administered once the patient has already seroconverted would affect replication (Fig. 4H). We find that nAbs can inhibit SARS-CoV-2 spread throughout the organoid, when compared to a nAb control targeting West Nile virus, highlighting the utility of our system to mimic *in vivo* situations (Fig 4I). Therefore, these antibodies may represent biologics to block SARS-CoV-2 infection and intercellular transmission to treat COVID-19 disease^31–32^. Moreover, our work highlights the utility of synthetic human lung buds to test candidate COVID-19 therapeutics in a high-throughput manner.

**Figure 4:**
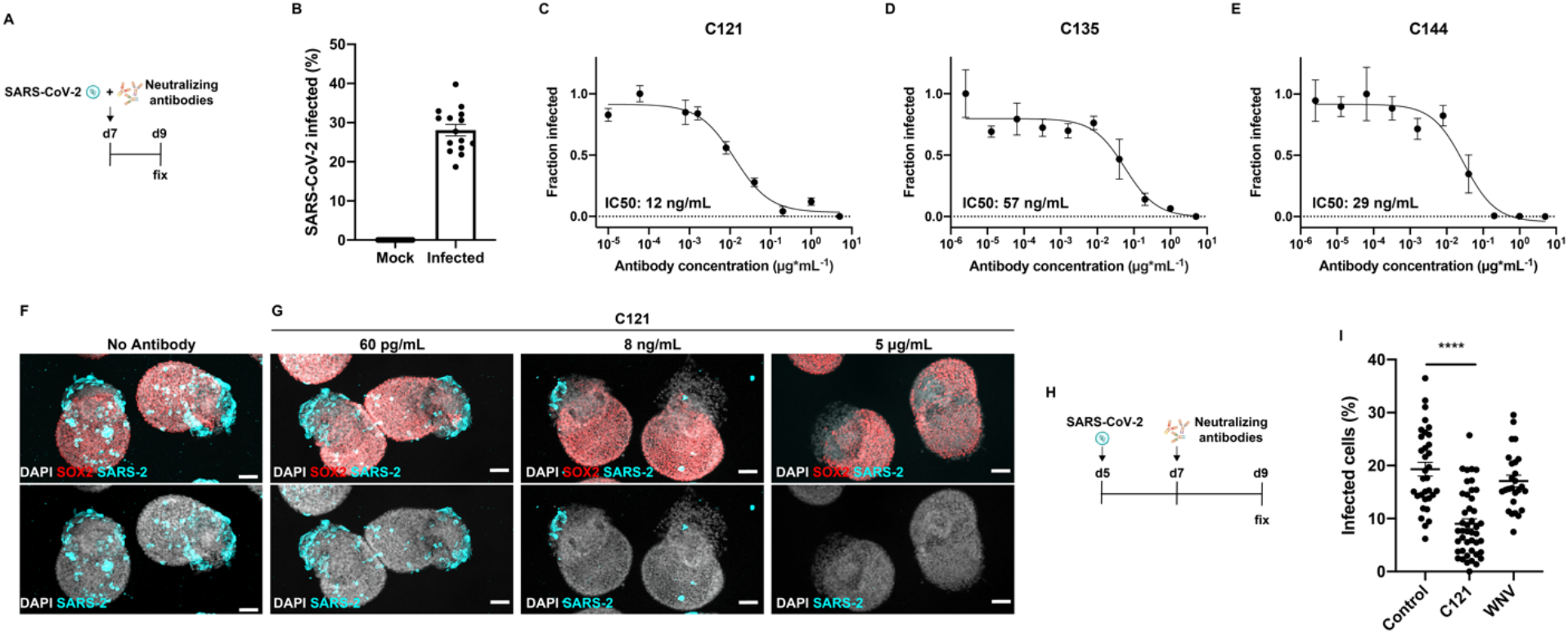
Evaluation of antibodies that neutralize SARS-CoV-2 infection. A) Experimental scheme for the high-throughput-based testing of SARS-CoV-2-neutralizing antibodies. B) Quantification of infected cells in control mock and SARS-CoV-2 infected synthetic lung buds in a high-throughput platform. C-E) Neutralization curves of SARS-CoV-2 infection in synthetic lung buds at increasing doses of three human antibodies. Data normalized to maximum (1) and minimum (0) infection levels. F-G) Representative images of synthetic lungs infected with SARS-CoV-2 and treated with varying doses of C121 human antibody (scale bar: 50μm). H) Experimental scheme to test the effect on neutralizing antibodies on intercellular transmission. Synthetic lung buds were infected with SARS-CoV-2 at day 5 of lung bud formation and treated with neutralizing antibodies 48 hours later. I) Proportion of infected cells in synthetic human lungs treated with C121 or a West Nile virus (WNV)-specific antibody from day 7 to day 9. (****p<0.0001, Dunnett’s multiple comparison test).

## Conclusion

Our work highlights the self-organizing capabilities of lung progenitors when they are cultured on confined geometries. Our synthetic human lung bud model recapitulates the earliest events in fetal lung development which involves the proximo-distal coordination of alveolar and airway cellular differentiation and region-specific tissue morphogenetic events^20–21^. It also offers key advantages to current *in vitro* models as it displays *in vivo*-like tissue organization and complexity as well as it provides inexhaustible access to disease-relevant lung tissue. Our work delineates novel cell types and cellular ontogeny in fetal human lung buds at the tissue and single cell level. This will allow for the interrogation of the molecular and genetic mechanisms orchestrating early human lung differentiation and morphogenesis, in both normal and disease states in which these processes may go awry such as lung cancer and other diseases. Our work also provides fast and scalable access to lung tissue for regenerative medicine and modeling lung diseases.

This study also highlights the potential of using synthetic lungs to identify novel therapeutics that block infection by SARS-CoV-2, endemic coronaviruses as well as other respiratory viruses. Compared to 3D organoid models where tracking virus-induced cellular responses is challenging due to inter-organoid variability in cell type complexity and composition^18^, our platform allows for the individual tracking of many genetically-matched reproducible organoids at a time to gain a quantitative understanding of SARS-CoV-2-induced lung pathology. We have established a highly quantitative platform to assess differential cell type-dependent susceptibilities to infection and cytopathology as well as to identify key target cell types that are otherwise inaccessible for experimentation *in vivo*. Whether our lung model recapitulates infection in fetal and adult tissue *in vivo* warrants further investigation. As we have detected tropism of other endemic coronaviruses (Extended Data Fig. 6), our platform will help gain an understanding of the cellular mechanisms that lead to the increased pathogenic effects and infectivity of SARS-CoV-2 compared to other related viruses. Importantly, as our approach is also amenable for high-throughput genetic analysis, it will unveil the complement of host factors that control lung infection which may represent therapeutic targets for COVID-19 and other emerging diseases as well. We expect our synthetic human lung model to be transformative for the elucidation of the molecular basis of respiratory infections and lung diseases for which there are currently no treatments.

## Methods

### Maintenance of hPSCs

RUES2 (NIHhESC-09-0013), RUES2-GLR^33^ and HUES8 iCas9 (NIHhESC-09-0021) hPSC lines were used in this study and maintained in HUESM (DMEM supplemented with 20% knockout serum replacement, 1×B27 supplement without vitamin A, 0.1 mM non-essential amino acids, 2 mM GlutaMax and 0.1 mM 2-mercaptoethanol) conditioned by mouse embryonic fibroblasts (MEF-CM) and supplemented with 20 ng*ml^−1^ basic fibroblast growth factor (bFGF). The cells were grown at 37 °C and 5% CO2 on tissue culture dishes that were coated with Geltrex (Life Technologies) solution.

### Generation of lung buds in micropatterns

hPSCs were first differentiated into definitive endoderm using the STEMDiff Definitive Endoderm Kit (Stem Cell Technologies), with 1 day addition of supplement CJ and MR and 2 days addition of supplement CJ only. Upon endoderm induction, cells were washed once with PBS^−/−^ (Gibco) and dissociated with Accutase (Stem Cell Technologies) for 7 mins. Cells were then dissociated with a pipette to ensure a single cell suspension and diluted in 4 times complete serum-free differentiation medium (cSFDM) containing DMEM/F12 (Gibco) with B27 Supplement with retinoic acid (Invitrogen, Waltham, MA), N2 Supplement (Invitrogen), 0.1% bovine serum albumin Fraction V (Invitrogen), ß-mercaptoethanol (Sigma), Glutamax (ThermoFisher), 50 μg*mL^−1^ ascorbic acid (Sigma), and normocin with supplements of 10 μm SB431542 (‘‘SB’’; Tocris), 2 μm Dorsomorphin (‘‘DS’’; Stemgent) and 10 μm ROCK inhibitor (Y-27632; Abcam). Cells were further diluted with the same medium and 5 × 10^5^ cells in 3.0 ml of medium were placed over a laminin-coated micropattern glass coverslips (CYTOOCHIPS Arena A, Arena 500A, Arena EMB A, Arena 225A)^22^ in a 35-mm tissue culture dish, left untouched for 10 mins and then incubated at 37 ° C. After 3 hours, the micropattern was washed once with PBS^+/+^, which was then replaced with cSFDM with 10 μm SB and 2 μm DM. 2 days later, the media was replaced with cSFDM with 10 μm SB and 2 μm DM. The next day, micropattern cultures were fed in lung induction medium (LIM) which contains cSFDM supplemented with 50 ng*mL^−1^ KGF (or indicated in each experiment; R&D systems), 10 ng*mL^−1^ BMP4 (R&D systems), 100 nM retinoic acid (“RA”; Sigma Aldrich) and 3 μM CHIR9902 (EMD Millipore). Micropattern cultures were fed every other day in LIM for 7 days or until tissues were collected for analysis.

To generate synthetic lung buds on 96 well plates, 4×10^4^ definitive endoderm cells were seeded on laminin-coated 96 well plates (CYTOOPlates, 200A), left untouched for 1 hour and then incubated at 37 ° C for 3 hours in cSFDM with 10 μm SB, 2 μm DM and 10 μm ROCK inhibitor. After this point, the protocol described above to generate lung buds on coverslips was followed.

### Immunostaining

Micropattern coverslips were fixed with 4% paraformal-dehyde (Electron Microscopy Sciences 15713) in warm medium for 30 min, rinsed three times with PBS^−/−^, and then blocked and permeabilized with 3% normal donkey serum (Jackson Immunoresearch 017-000-121) with 0.5% Triton X-100 (Sigma 93443) in PBS^−/−^ for 30 min. Micropatterns were incubated with primary antibodies for 1.5 h, washed three times in PBS^−/−^ for 5 min each, incubated with secondary antibodies conjugated with Alexa 488, Alexa 555, Alexa 594 or Alexa 647 (1:1,000 dilution, Molecular Probes), fluorescently-conjugated phalloidin (1:400; Life Technologies) and 10 ng*ml^−1^ of DAPI (Thermo Fisher Scientific D1306) for 30 min and then washed three times with PBS^−/−^. Coverslips were mounted on slides using ProLong Gold antifade mounting medium (Molecular Probes P36934).

The primary antibodies used were as follows: rabbit anti-SOX9 (Millipore; AB5535; 1:250), goat anti-SOX2 (R&D Systems; AF2018; 1:250), rabbit anti-NKX2.1 (Abcam; ab76013; 1:200), mouse anti-Acetylated Tubulin (Sigma; T7451; 1:1000), goat anti-TP63 (R&D Systems; BAF1916; 1:250), rabbit anti-proSPC (Seven Hills; WRAB-9337; 1:500), mouse anti-HOPX (Santa Cruz; sc-398703; 1:250), rabbit anti-nucleocapsid SARS-CoV-2 (GeneTex; GTX135357; 1;1000); rabbit anti-Active Caspase-3 (R&D Systems; AF835; 1:250); mouse anti-Mucin 5AC (Abcam; ab3649; 1:250); human anti-Spike SARS-CoV-2 (Robbiani et al., 2020; 1:1,000); anti-HNF-3BETA/FOXA2 (Neuromics; GT15186; 1:200); mouse J2 dsRNA (SCICONS; 1;1,000), phospho Histone H3 (Cell Signaling; 9706S; 1:200) and goat anti-SOX17 (R&D Systems; AF1924; 1:200). To detect infected cells for HCoV-229E, HCoV-OC43 and HCoV-NL63, a mouse monoclonal anti-dsRNA antibody (Scicons: catalog no. 10010500) was used under similar conditions.

### Imaging

All confocal images were acquired on a Zeiss Inverted LSM 780 laser scanning confocal microscope with a 10X, 20X or 25X oil-immersion objective. 96 well plates were imaged on a ImageXpress Micro with a 10X objective. Three-dimensional visualization and image processing was performed in ImageJ.

### Imaging analysis

Analysis of confocal Z-stacks was performed using CellProfiler v4.0.4 (https://cellprofiler.org/). DAPI staining was used to identify nuclei and cytoplasm. Cells were classified as positive for a given marker based on the identification of fluorescence signals in either nuclear (for nuclear markers such as SOX9 and SOX2) or cytoplasmic area (SARS-CoV-2 and aCASP3). The total percentage of cells positive for a given marker was calculated as the sum of the number of cells identified as positive for the marker in each Z section divided by the total number of cells identified.

96-well plates image analysis was performed using ImageJ. Briefly, nuclei were identified and set to a mask. The ROIs were then expanded, creating a new mask to interrogate cytoplasmic signals of virus infection in the SARS-CoV-2 images.

### Single cell RNA-sequencing analysis

Micropatterned coverslips with 225-μm diameter synthetic lung buds at day 7 of lung induction were dissociated with TrypLE Express (Gibco) for 10 min at 37 °C. After dissociation, the cells were washed three times in PBS^−/−^ (Gibco) with 0.04% BSA and strained through a Flowmi tip 40 μm strainer. Cell count and viability were determined on a Countess II Automated Cell Counter. Samples were loaded for capture with the Chromium System using the Single Cell 3′ v3 reagents (10X Genomics). Following cell capture and lysis, cDNA was synthesized and amplified according to the manufacturer’s instructions (10X Genomics). The resulting libraries were sequenced on the Nova-seq platform. The Cell Ranger (v.2.0.2) software pipeline was used to create FASTQ files which were aligned to the hg19 genome using default parameters. These data are available through the NCBI GEO accession number GSE163698.

Filtered gene expression matrices were generated using Cell Ranger and subsequently used for downstream analyses using Scanpy (v.1.6.0) (https://pypi.org/project/scanpy/). Data was filtered to have a minimum of 2000 and a maximum of 8000 detected genes per cell and genes were filtered to be expressed in at least 3 cells. Cells with over 10% mitochondrial genes were discarded. Clustering of cells was performed using the Leiden algorithm (resolution=0.4) and visualized using UMAP plots (n.neighbors = 50, PCA components= 50). Differential gene expression analysis was performed using the Wilcoxon rank-sum (Mann-Whitney-U) test to identify cluster markers.

### RNA velocity

Count matrix layered with spliced and unspliced annotations were obtained using Kallisto Bustools^34^. In brief, kb-python==0.24.4 was used to capture spliced and unspliced transcripts using -lamanno function with the GRCm38 human genome. The Kallisto unfiltered count matrix was imported in Scanpy and further processed for scVelo (v0.2.2) analysis^26–27^. UMAP coordinates and clusters annotation from the Scanpy clustering were used.

### SARS-CoV-2 infection and transmission analysis

SARS-CoV-2 (strain: USA-WA1/2020) and HCoV-NL63 were obtained from BEI Resources (NR-52281 and NR-470). HCoV-OC43 was obtained from ZeptoMetrix (cat. #0810024CF) and HCoV-229E was generously provided by Volker Thiel (University of Bern). All viruses were amplified at 33 °C in Huh-7.5 cells to generate a P1 stock. To generate working stocks, Huh-7.5 cells were infected at a multiplicity of infection (MOI) of 0.01 plaque forming unit (PFU)/cell (SARS-CoV-2, HCoV-NL63, HCoV-OC43) and 0.1 PFU/cell (HCoV-229E) and incubated at 33 °C until virus-induced CPE was observed. Supernatants were subsequently harvested, clarified by centrifugation (3,000 g × 10 min) at 4 dpi (HCoV-229E), 6 dpi (SARS-CoV-2, HCoV-OC43) and 10 dpi (HCoV-NL63), and aliquots stored at −80 °C.

Viral titers were measured on Huh-7.5 cells by standard plaque assay. Briefly, 500 μL of serial 10-fold virus dilutions in Opti-MEM were used to infect 4 × 105 cells seeded the day prior into wells of a 6-well plate. After 90 min adsorption, the virus inoculum was removed, and cells were overlaid with DMEM containing 10% FBS with 1.2% microcrystalline cellulose (Avicel). Cells were incubated for 4 days (HCoV-229E), 5 days (SARS-CoV-2, HCoV-OC43) and 6 days (HCoV-NL63) at 33 °C, followed by fixation with 7% formaldehyde and crystal violet staining for plaque enumeration. All SARS-CoV-2 experiments were performed in a biosafety level 3 laboratory.

### Antibody treatments

Neutralization assays were performed as previously described^30^. Briefly, antibodies were serially diluted in LIM, mixed with a constant amount of SARS-CoV-2 (2 × 10^5^ PFU for the larger chips, and 5 × 10^4^ PFU in the 96-well plate assays) and incubated for 60 min at 37 °C. The antibody– virus mix was then added to the 96-well plates or microchip containing synthetic human lungs.

### Statistical analysis

Statistical analysis was performed using unpaired two-sided t tests, unless stated otherwise. For all the experiments included in this study, three or more biological replicates were performed using stage-matched controls as a reference. No statistical analysis was used to predetermine sample size and no data were excluded.

## Figure legends

**Extended Data Figure 1: Induction of NKX2.1+ lung progenitors on confined geometries.** A) Monolayer differentiation of SOX17+ endoderm progenitors pre-seeding. B) SOX17+ endoderm progenitors on confined geometry 3 hours post-seeding. C-E) Induction of NKX2.1+ multipotent lung and SOX2+ airway progenitors upon modulation of BMP4, KGF or KGF+BMP4. (scale bar: 50 μm). F) Efficient induction of NKX2.1+ multipotent lung and SOX2+ airway progenitors in colonies of varying sizes. (scale bar: 100 μm)

**Extended Data Figure 2: KGF- and BMP4-dependent induction of NKX2.1+ lung progenitors.** A) Low magnification images of epithelial buds containing NKX2.1+ lung progenitors grown on confined geometries of 225 μm diameter at varying doses of KGF. Epithelial structures can be identified with F-actin staining. B) Induction of NKX2.1+ lung progenitors on confined geometries of 500 μm diameter at varying doses of BMP4. C) Proportion of NKX2.1+ cells at varying doses of BMP4. (**p<0.01, ****p<0.0001, Dunnett’s multiple comparison test)

**Extended Data Figure 3: Expression of cell type markers in 500 μm colonies after lung progenitor induction.** A) Expression of SOX2 and SOX9 in non-overlapping tissue domains. B-E) Identification of AcTub+ multiciliated cells (B), P63+ airway basal stem cells (C), MUC5AC+ goblet cells (D) and proSFTPC+ type 2 pneumocytes (E). (scale bar: 50μm)

**Extended Data Figure 4: Single-cell gene expression analysis of synthetic lung buds.** A) Heatmap of top 10 differentially expressed genes for each cluster identified in synthetic lung buds. z-score normalized expression values are shown. B) Violin plots of the number of genes (n_genes) and percentage of mitochondrial genes (pct_counts_mt) for each cluster identified in synthetic lung buds. C) Cluster-level gene expression Pearson correlation analysis of clusters identified in synthetic lung buds. z-score normalized correlation values are shown. D) UMAP expression plots of SOX9, SOX2 and lumican (LUM). E) Gene expression trackplots of cell type-specific markers. Each peak represents a single cell and its height denotes the expression level of each gene.

**Extended Data Figure 5: Single cell gene expression analysis of synthetic lung buds.** A) Heatmap of scaled gene expression levels of alveolar candidate markers. B-D) Pseudotime-aligned gene expression heatmap of top 300 genes differentially expressed along the global differentiation trajectory (B) as well as AT1/2 to AT1/2s (C) and AT1/2 to AT1 (D) trajectories identified by RNA velocity. Color bar on top of the heatmap corresponds to the cluster classification in Fig. 2G for each cell aligned along differentiation pseudotime. E) UMAP plots and classification of cell types in synthetic human lung buds. F) UMAP expression plots of ACE2, TMPRSS2 and FURIN. C) Scaled expression value of entry factors for each of the identified clusters in synthetic human lung buds.

**Extended Data Figure 6: Infection of synthetic lung buds by endemic coronaviruses.** A-D) Synthetic lung buds infected with endemic coronaviruses HCoV-229E (B), HCoV-OC43 (C) and HCoV-NL63 (D). Infected cells were identified by staining with J2 antibody detecting dsRNA.

**Extended Data Figure 7: Infection levels in alveolar and airway cells across developmental stages.** A-C) Percentage of total (A), SOX9+ (B) and SOX2+ (C) cells infected by SARS-CoV-2 at multiple stages of lung bud formation.

## Acknowledgements

We would like to thank C. Zhao and the Genomics Resource Center at the Rockefeller University for help and advice regarding single-cell RNA sequencing as well as A. North and the Bio-Imaging Resource Center for advice regarding imaging analysis. We thank members of the Brivanlou and Rice laboratories for critical discussions and comments on the manuscript. This work was supported by the Pershing Square Foundation, NIH grants P01AI138398-S1, 2U19AI111825, and R01AI091707-10S1, a George Mason University Fast Grant, the BAWD Foundation, the G. Harold and Leila Y. Mathers Charitable Foundation, and private funding from the Rockefeller University.

## Author Contributions

E.A.R. and A.H.B conceived and designed the lung organoid platform. E.A.R., B.R., H.H., C.M.R. and A.H.B. designed experiments with SARS-CoV-2. E.A.R. and B.R. executed and analyzed the experiments. E.A.R. and R.D.S performed the single cell gene expression analysis. E.A.R, B.R., C.M.R. and A.H.B. wrote the manuscript with input from all authors.

## Materials & Correspondence

The datasets generated during and/or analyzed during the current study are available from the corresponding authors C.M.R. and A.H.B. on reasonable request.

