## SupplementaryData for "Self-organized stem cell-derived human lung buds with proximo-distal patterning and novel targets of SARS-CoV-2"

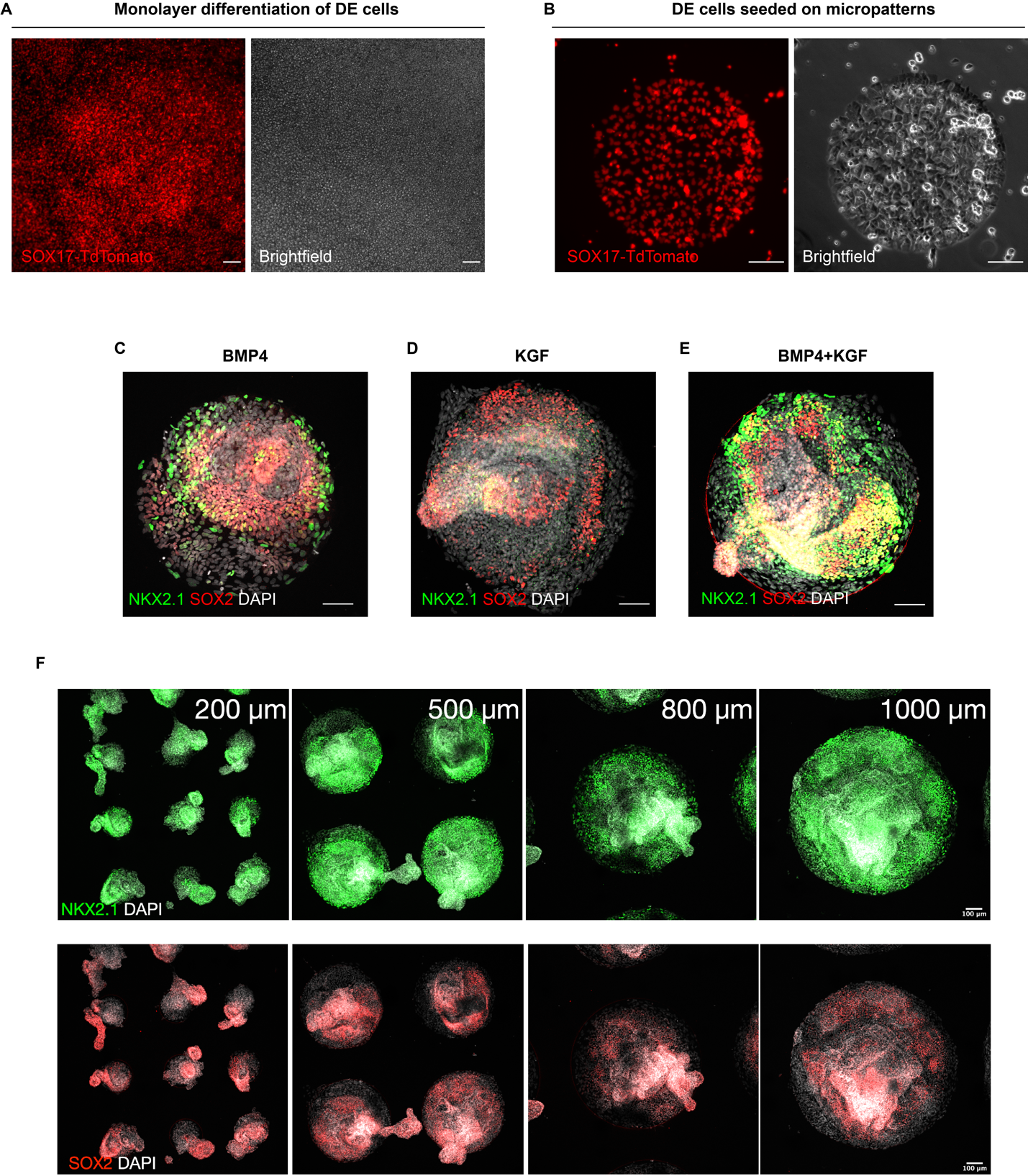

**Extended Data Figure 1: Induction of NKX2.1+ lung progenitors on confined geometries.** A) Monolayer differentiation of SOX17+ endoderm progenitors pre-seeding. B) SOX17+ endoderm progenitors on confined geometry 3 hours post-seeding. C-E) Induction of NKX2.1+ multipotent lung and SOX2+ airway progenitors upon modulation of BMP4, KGF or KGF+BMP4. (scale bar: 50 µm). F) Efficient induction of NKX2.1+ multipotent lung and SOX2+ airway progenitors in colonies of varying sizes. (scale bar: 100 µm)

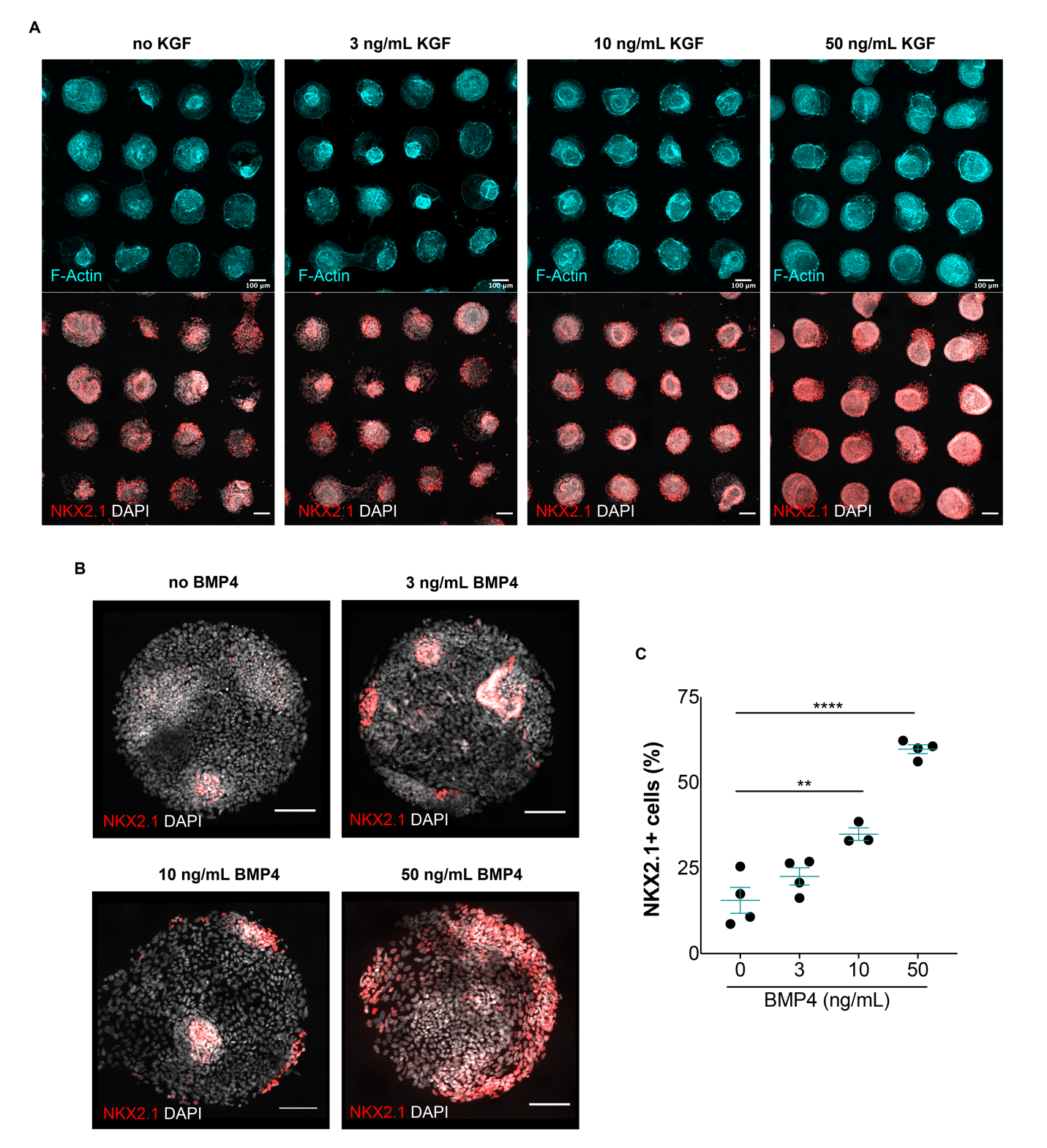

**Extended Data Figure 2: KGF- and BMP4-dependent induction of NKX2.1+ lung progenitors.** A) Low magnification images of epithelial buds containing NKX2.1+ lung progenitors grown on confined geometries of 225 µm diameter at varying doses of KGF. Epithelial structures can be identified with F-actin staining. B) Induction of NKX2.1+ lung progenitors on confined geometries of 500 µm diameter at varying doses of BMP4. C) Proportion of NKX2.1+ cells at varying doses of BMP4. (**p<0.01, ****p<0.0001, Dunnett’s multiple comparison test)

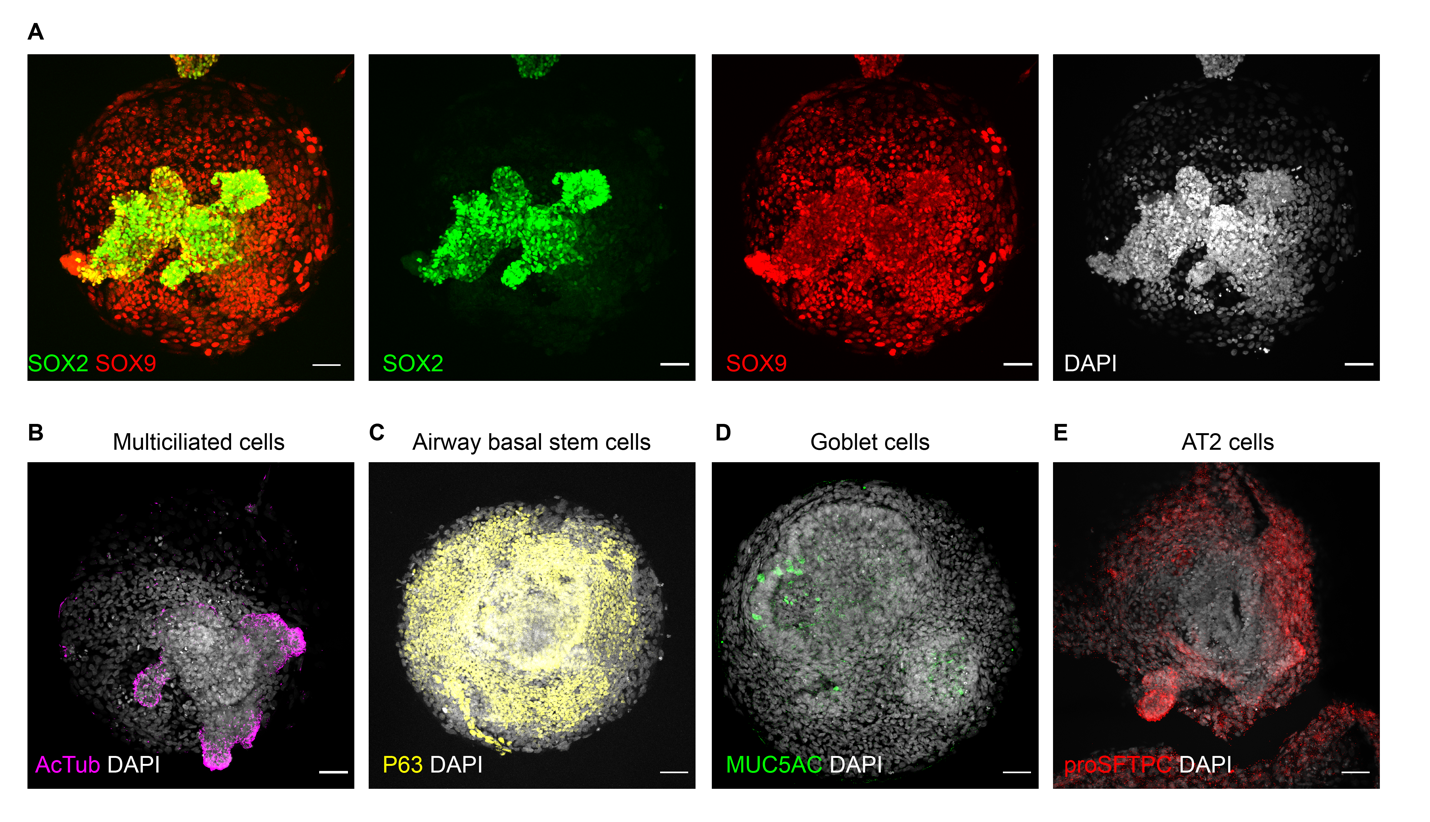

**Extended Data Figure 3: Expression of cell type markers in 500 µm colonies after lung progenitor induction.** A) Expression of SOX2 and SOX9 in non-overlapping tissue domains. B-E) Identification of AcTub+ multiciliated cells (B), P63+ airway basal stem cells (C), MUC5AC+ goblet cells (D) and proSFTPC+ type 2 pneumocytes (E). (scale bar: 50µm)

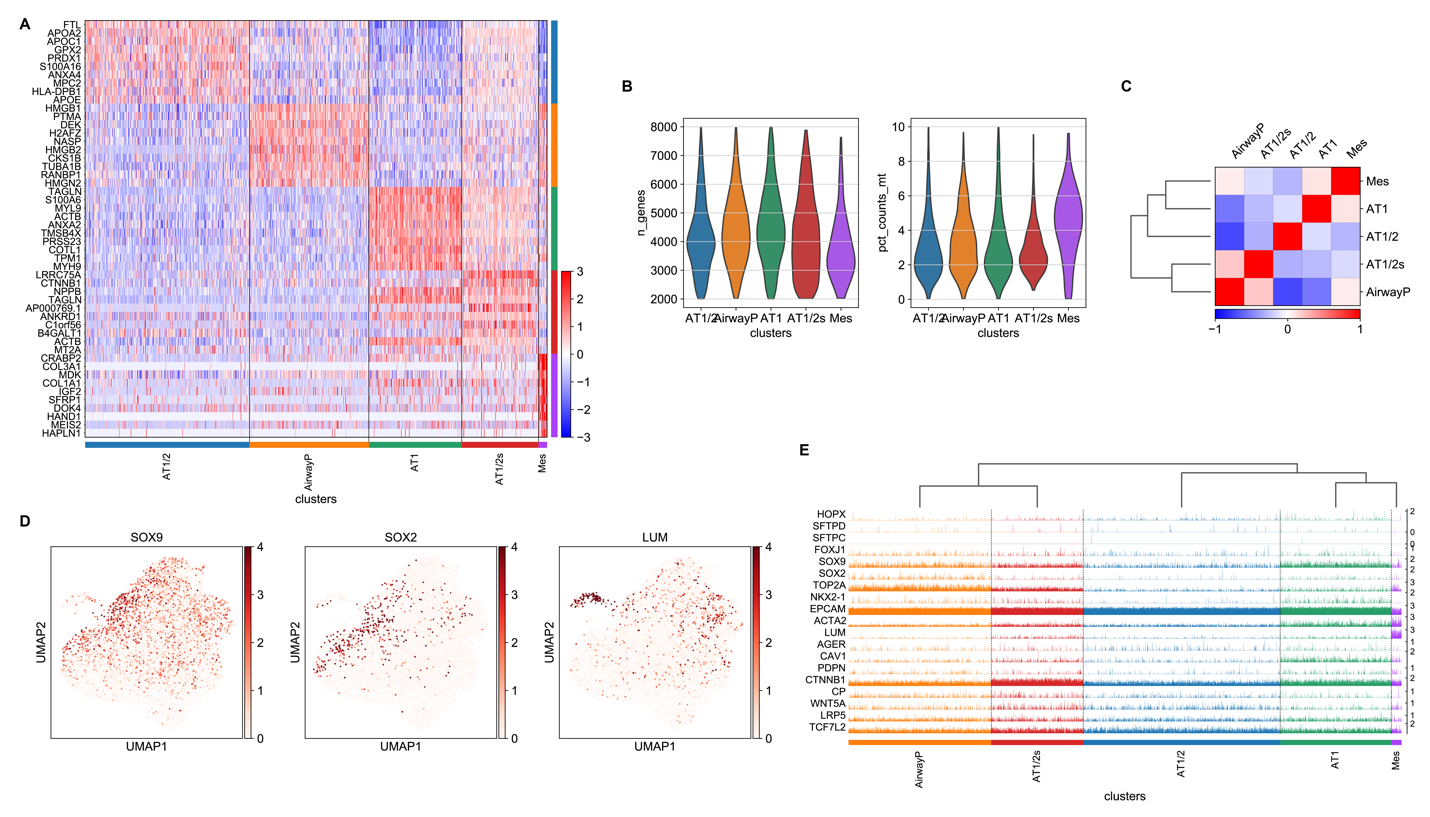

**Extended Data Figure 4: Single-cell gene expression analysis of synthetic lung buds.** A) Heatmap of top 10 differentially expressed genes for each cluster identified in synthetic lung buds. z-score normalized expression values are shown. B) Violin plots of the number of genes (n_genes) and percentage of mitochondrial genes (pct_counts_mt) for each cluster identified in synthetic lung buds. C) Cluster-level gene expression Pearson correlation analysis of clusters identified in synthetic lung buds. z-score normalized correlation values are shown. D) UMAP expression plots of SOX9, SOX2 and lumican (LUM). E) Gene expression trackplots of cell type-specific markers. Each peak represents a single cell and its height denotes the expression level of each gene.

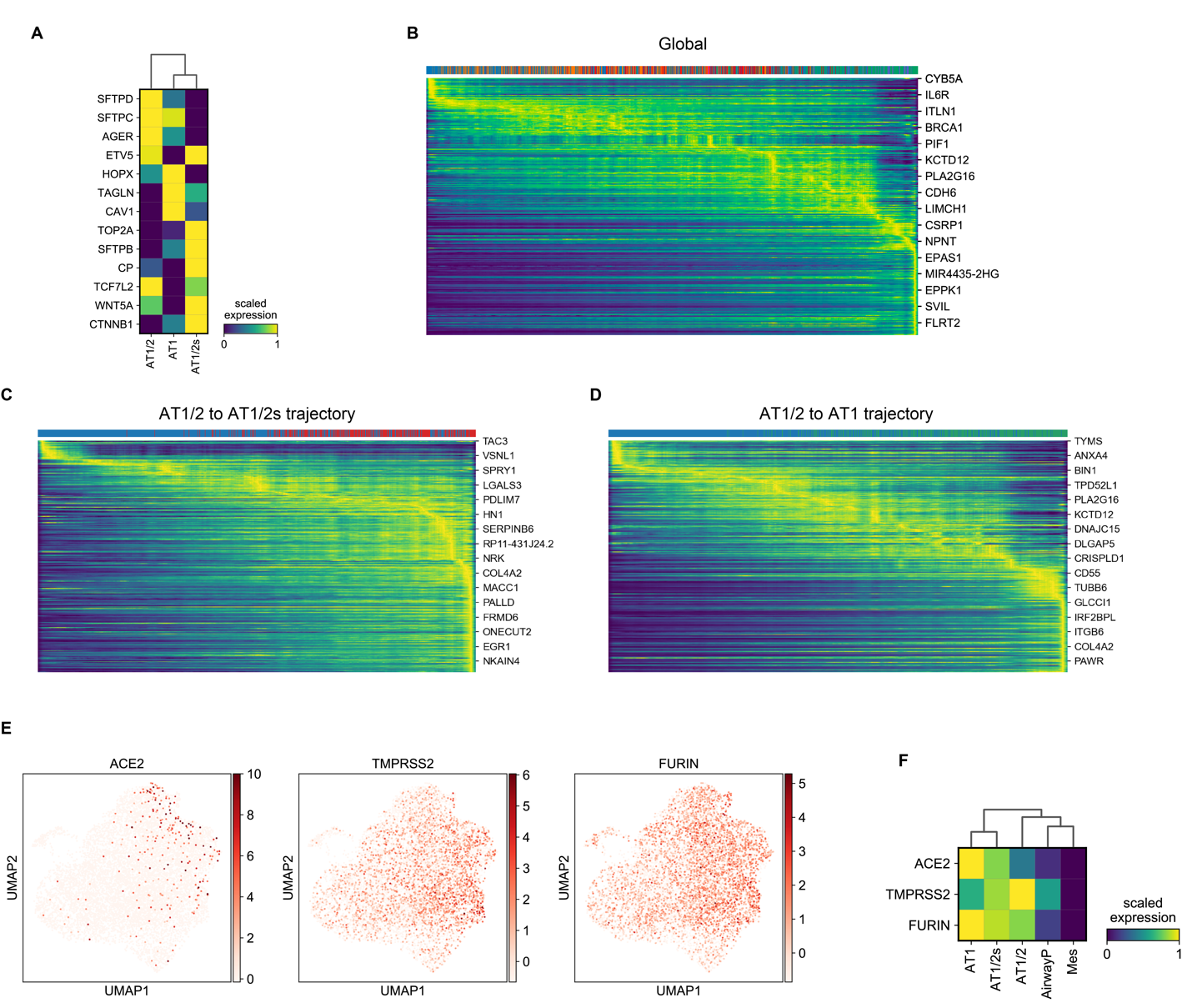

**Extended Data Figure 5: Single cell gene expression analysis of synthetic lung buds.** A) Heatmap of scaled gene expression levels of alveolar candidate markers. B-D) Pseudotime-aligned gene expression heatmap of top 300 genes differentially expressed along the global differentiation trajectory (B) as well as AT1/2 to AT1/2s (C) and AT1/2 to AT1 (D) trajectories identified by RNA velocity. Color bar on top of the heatmap corresponds to the cluster classification in Fig. 2G for each cell aligned along differentiation pseudotime. E) UMAP plots and classification of cell types in synthetic human lung buds. F) UMAP expression plots of ACE2, TMPRSS2 and FURIN. C) Scaled expression value of entry factors for each of the identified clusters in synthetic human lung buds.

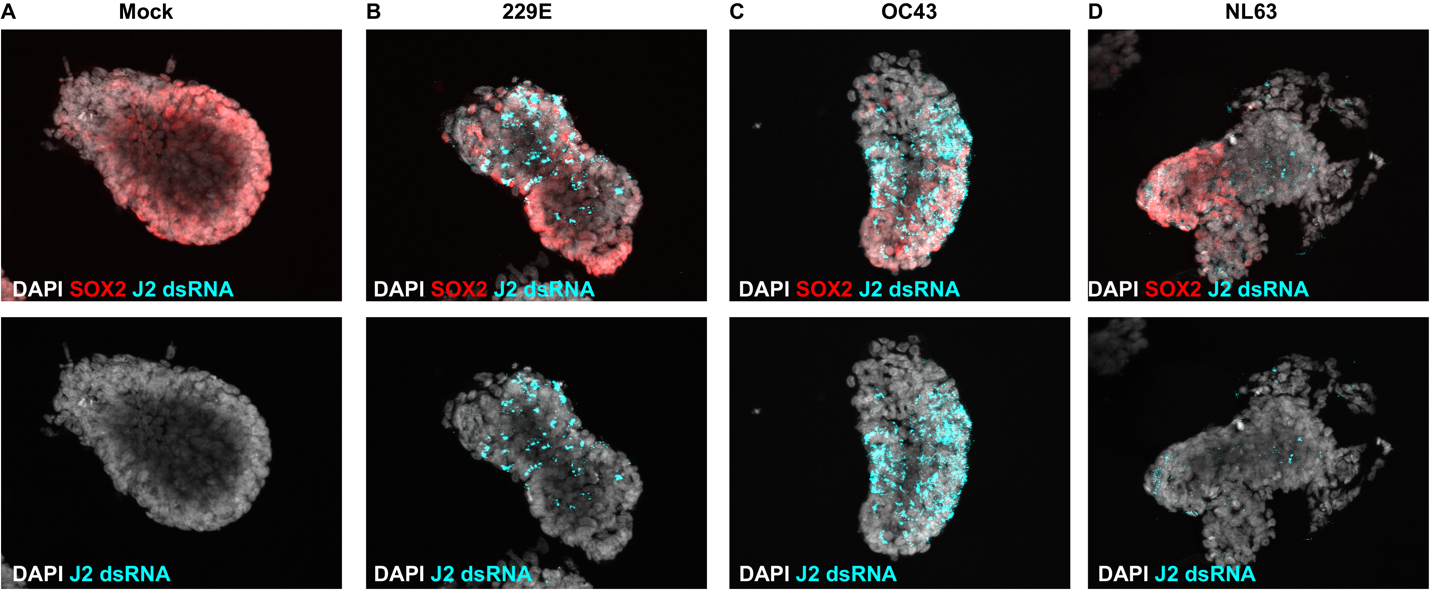

**Extended Data Figure 6: Infection of synthetic lung buds by endemic coronaviruses.** A-D) Synthetic lung buds infected with endemic coronaviruses HCoV-229E (B), HCoV-OC43 (C) and HCoV-NL63 (D). Infected cells were identified by staining with J2 antibody detecting dsRNA.

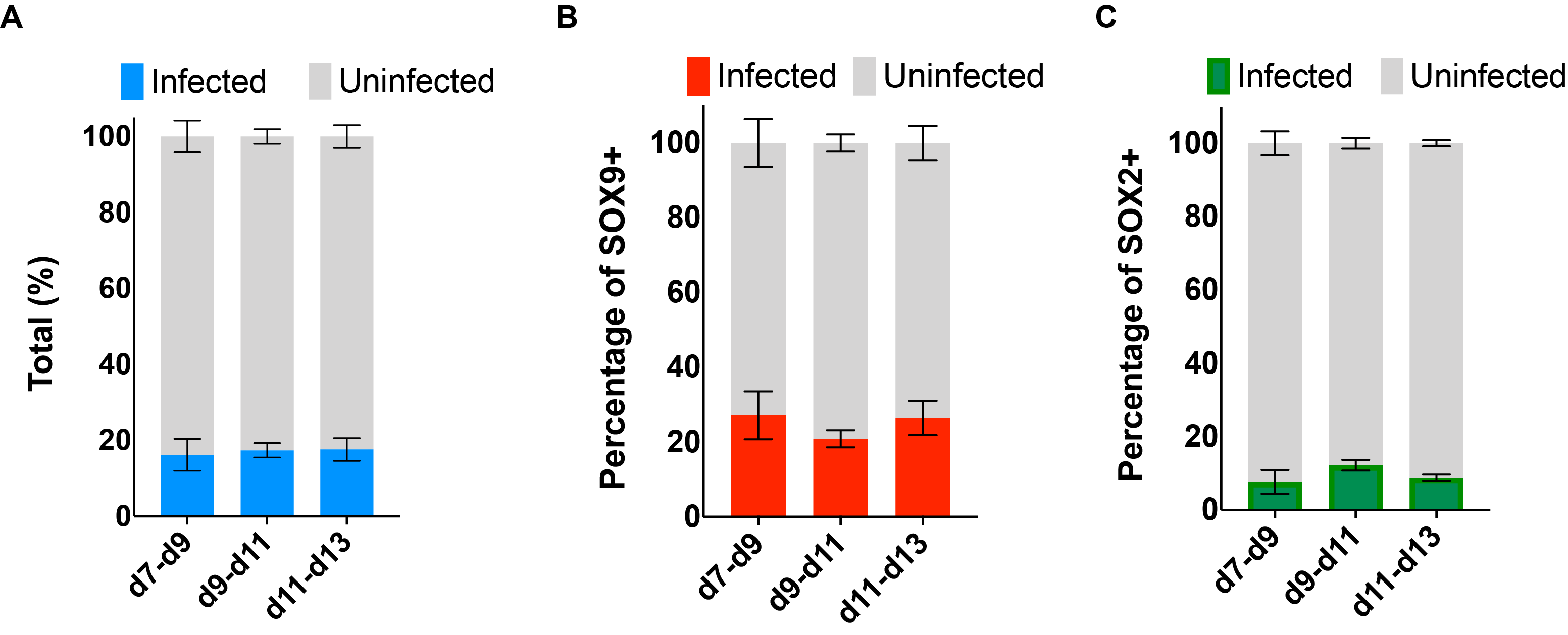

**Extended Data Figure 7: Infection levels in alveolar and airway cells across developmental stages.** A-C) Percentage of total (A), SOX9+ (B) and SOX2+ (C) cells infected by SARS-CoV-2 at multiple stages of lung bud formation.
